# Geometry representations along visual pathways in human spatial navigation

**DOI:** 10.1101/2024.08.04.605402

**Authors:** Taiping Zeng, Ming Bo Cai

## Abstract

The representation of geometric structures of one’s surroundings is key to self-localization during human spatial navigation. However, the spatial organization of geometry representation in the visual system has not been fully characterized. By modeling the synchronized brain activity from participants watching navigational videos from identical realistic virtual environments under different weather and lighting conditions, we found a compact representation of egocentric 3D scene geometric structures is present in a widespread network of brain regions. These regions carrying geometry-related signals form three parallel pathways, which we collectively refer to as “geometry visual pathways”, starting from the primary visual cortex: the dorsal and medial pathways end in the intraparietal areas, while the ventral pathway arrives at the hippocampus via the parahippocampal gyrus. The synchronized neural activity in the identified geometry visual pathways allows for reliable decoding of the 3D scene geometric structures. Furthermore, road types, a more abstract representation of the route geometry, are encoded in overlapping pathways, distinctly absent from early visual cortex (V1, V2, V3). In addition to complementing the classical “what” and “where” dichotomy, the identified brain-wide geometry visual pathways narrow a critical gap in understanding how the brain constructs cognitive maps from visual inputs.

## Introduction

Spatial navigation is a fundamental cognitive ability that enables humans to move purposefully through complex environments, underlying countless daily activities from way finding to precise object-directed actions. To localize ourselves during navigation, the brain needs to form and refer to cognitive maps [1, 2], a mental representation of the surrounding space. Evidence suggests that the brain maintains both a metric-like and a graph-like cognitive map of space [3] in the hippocampus and surrounding areas [4, 5]. The visual system is considered to contain separate “what” and “where” pathways that both support spatial navigation. Particularly, the “where” visual pathway is suggested to generate the geometry representations required to build cognitive maps [6].

To form a metric-like cognitive map, the brain needs to infer the layout of the surrounding environment, which in turn requires the detection of boundaries in the environment. Neurons encoding for egocentric boundary have been found in dorsomedial striatum [7], lateral entorhinal cortex [8], and retrosplenial cortex [9]. Many other types of boundary-related neurons encoding for allocentric geometric information have also been identified [10], including boundary vector cells (BVCs) [11], border cells [12], boundary-off cells [13], and all-boundary cells, including “annulus” and “bulls-eye” cells [14] in multiple brain regions. We have previously theoretically hypothesized that another set of “geometry cells” [15] can further integrate boundary information in the postrhinal cortex [16] to infer the spatial layout of local scenes. While metric-like representations enable accurate navigation within local environments, the brain also constructs abstract graph-like maps of larger global environments to support efficient route planning [5]. In a graph-like map of a complex surrounding environment, local spatial layouts can be abstracted into reachable nodes and connecting edges. For example, in urban cities, route planning can be facilitated by representing the road network as an abstract graph in which intersections (T-junctions and crosses) serve as nodes that are connected by edges representing unbranched road segments. These road structure types (unbranched roads, turns, T-junctions, and crosses) can be considered as spatial layouts inferred based on environmental boundaries. However, despite the advances of identifying boundary-related neurons, it remains incompletely understood how 3D scene geometry and abstract navigational affordances are jointly represented across the visual pathways, especially under naturalistic continuous navigation.

Human neuroimaging studies have shown that the occipital place area (OPA) encodes the 3D surface layout of scenes [17, 18], while the parahippocampal place area (PPA) and retrosplenial cortex (RSC) contribute to scene identity and navigation [19, 6]. OPA also represents navigational affordances such as potential paths [20], providing a direct link between visual geometry and behavior. However, most of these studies used static images or short clips with relatively simplified models of scene layout. Here, we extend this line of work by analyzing brain activity during continuous naturalistic navigation videos. Using a deep generative model, we extract compact latent features representing scene depth structures from visual inputs that generalize across weather and lighting conditions, and annotated the types of road structures at the camera’s locations, which allow us to examine how the brain encodes these geometric representations during continuous navigation.

The “where” visual pathway is thought to be responsible for processing the spatial properties of objects and scenes to form 3D scene geometry representations (e.g., depth surfaces and layouts) [21, 22]. 2D images falling onto the retina are converted into egocentric 3D scene structures [17], which are used to infer the boundaries of scenes [23]. These geometry representations can further support more abstract navigational affordances, such as road topology (e.g., unbranched roads, turns, junctions), which are building blocks of cognitive maps with metric-like and graph-like features. From a feed-forward perspective, representations of 3D scene geometry and road structure should emerge sequentially from the early visual cortex to the hippocampus, but another perspective of a hierarchical generative model suggests these intermediate-level variables are distributed across a wide range of brain regions due to bidirectional information flow constraining perceptions at multiple levels [24, 25]. Here, we use a novel fMRI paradigm to identify the distribution of representations of coarse geometry of scene depth and abstract road types in the human visual system.

Our results revealed that a large network of geometry-selective brain regions forms a continuous map from the occipital lobe to the parietal lobe along a dorsal pathway and a medial pathway, as well as a ventral pathway leading to the hippocampus. Evidence not only confirms the involvement of the dorsal visual pathway for encoding 3D scene geometry, but also reveals overlapping regions that additionally represent navigational affordances. Taken together, these findings suggest multiple streams of brain regions that carry geometric information co-exist, which we jointly refer to as the ‘geometry visual pathways’.

## Results

To better understand geometry representations along visual pathways during human spatial navigation, we designed a novel fMRI experiment in which all participants watched first-person-view videos from a front-facing camera on a car navigating in the same set of realistic virtual environments but under different weather and lighting conditions (Figure 1). Then, using shared response modeling [26], a hyperalignment [27] algorithm, we projected fMRI responses from each participant into a shared low-dimensional space in which neural signals are synchronized, thus invariant to the differences in texture and color induced by weather or lighting conditions. We investigated the representation of global 3D geometry of scenes captured by a deep neural network, which is tasked to perform depth perception while using a low-dimensional representation to generate the depth field of many scenes (Figure 2). By calculating the variance of the synchronized neural signals in each brain region explained by the time courses of the abstract geometry representation in the neural network, we found that the videos evoked activity related to 3D geometry structure in a large network of brain regions, forming a continuous map from the occipital lobe to the parietal lobe along dorsal and medial visual pathways and a ventral pathway leading to the hippocampus, which we describe jointly as “geometry visual pathways” (Figure 3). Remarkably, abstract geometry representations, such as road structures, are mainly located in the higher-level visual regions but not in the early visual cortex (Figure 4). The geometric structures can be reconstructed from the neural signals in brain regions of the identified geometry visual pathways (Figure 5). Integrating other available evidence, we illustrate the connecting diagram of the geometry visual pathway along cortical streams in humans (Figure 6b).

**Figure 1.**
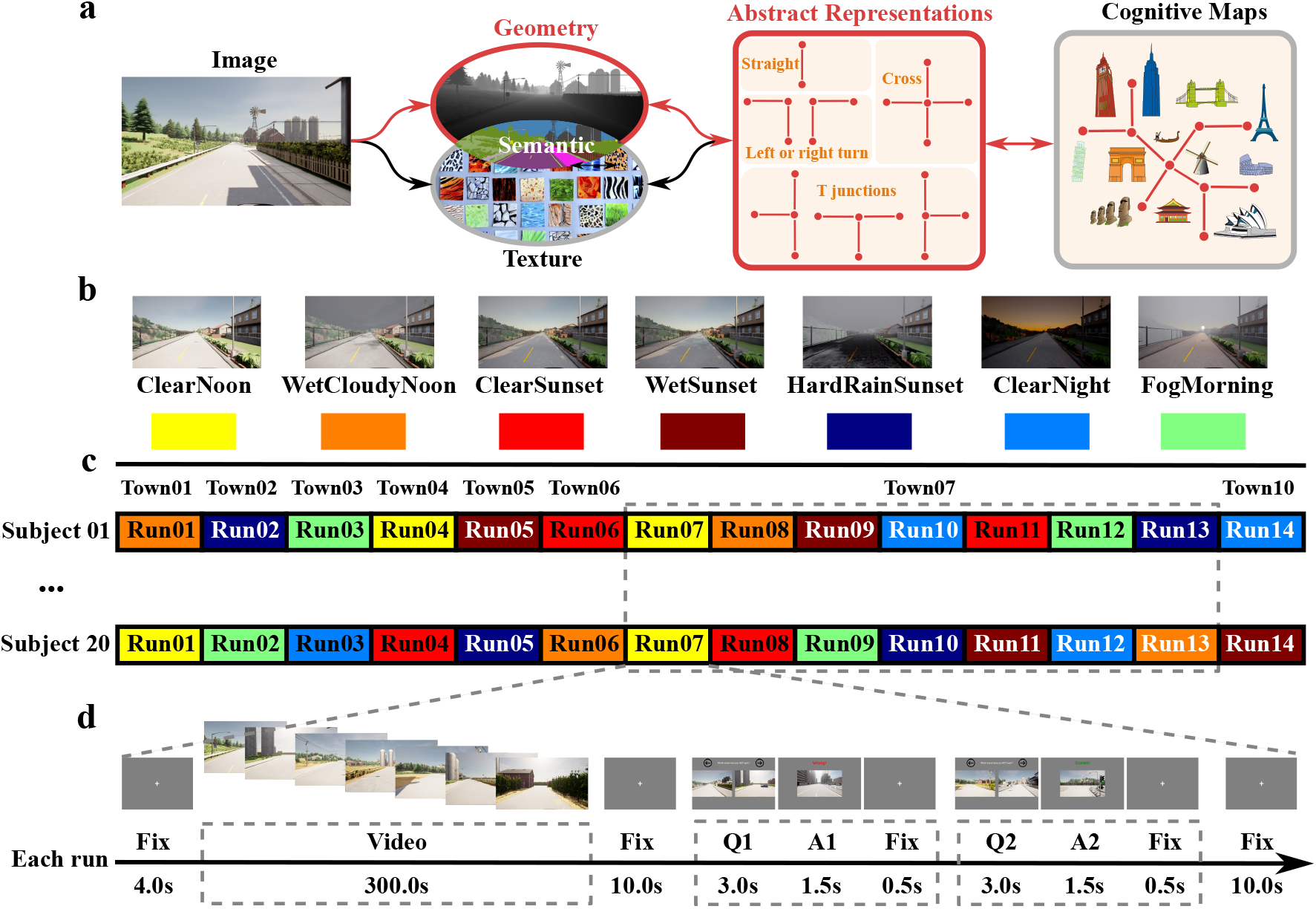
The fMRI experiment of geometry representations. (**a**) **The hypothetical framework of geometry representations in cognitive map formation**. The brain infers 3D geometry and texture information from the visual input. Semantic representation requires integration both geometry and texture information. They are used to further form the abstract representation of local road types, which is in turn required to construct cognitive maps. (**b**) **Example variation of weather and lighting conditions for the same scene**. Videos for each town were rendered in seven typical weather and lighting conditions, labeled by different colors. (**c**) **Experiment structure**. Each participant underwent 14 runs of fMRI scan. The weather and lighting condition were pseudo-randomly assigned across runs and participants. For Town07 which has more diverse environmental features, seven weather and lighting conditions were all viewed by each participant. (**d**) **Timeline of a single run**. Each run included watching a five-minute video and two memory tests that asked participants to report which of the two displayed scenes was not seen in the previous movie by pressing one of two buttons. The movie and questions were separated by fixation periods.

**Figure 2.**
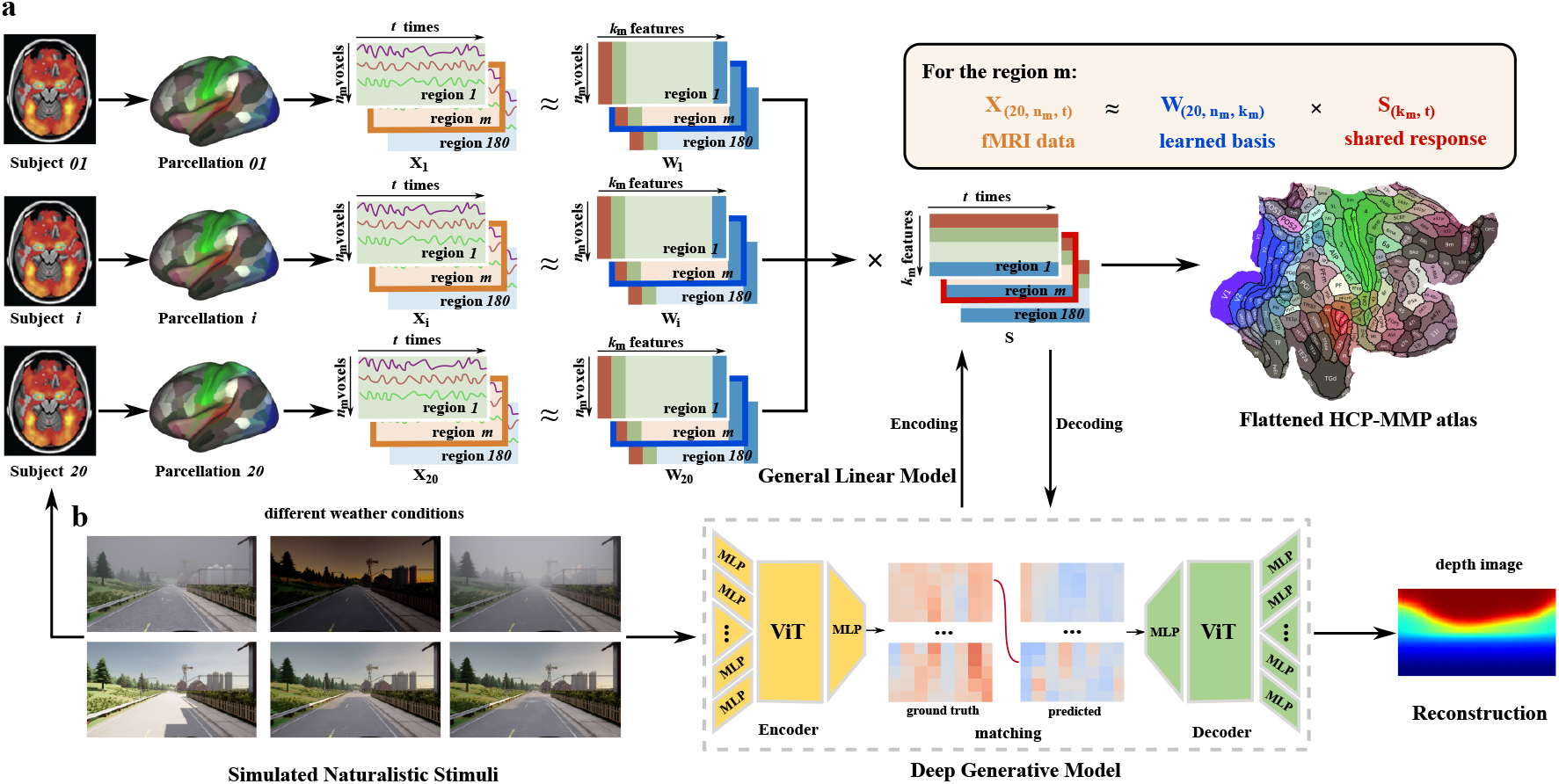
Schematic diagram of the model. (**a**) **Region-based shared response modeling**. The navigational videos watched by different participants during fMRI scans shared the same time courses of geometric structures, but had random weather and lighting conditions assigned to each participant. Each participant’s brain image was parcellated into 180 pairs of regions using the HCP-MMP atlas. Participant *i*’s data *X* in a region *m* are projected into a shared functional space *S* to capture geometry representations using shared response modeling: *X*_*mi*_ = *W*_*mi*_*S*_*m*_ + *ϵ* _*mi*_, where *X*_*mi*_, *W*_*mi*_ (learned spatial weights), and *ϵ*_*mi*_ (the residuals, not shown) are participant-specific, while *S*_*m*_ is shared across participants. (**b**) **Deep generative model**. A low-dimensional latent space is learned by a deep generative model (a vision transformer-based variational autoencoder using RGB as inputs and depth as outputs). A general linear model was fitted to explain variance in the shared neural responses using the time courses of the vectors in the latent space for the RGB images. A decoding model was trained to linearly predict the HRF-convolved latent vectors of the deep network from the shared neural responses, and reconstruct depth structure by identifying the ground truth latent vector across all scenes that best matches the decoded latent vector based on their correlation.

**Figure 3.**
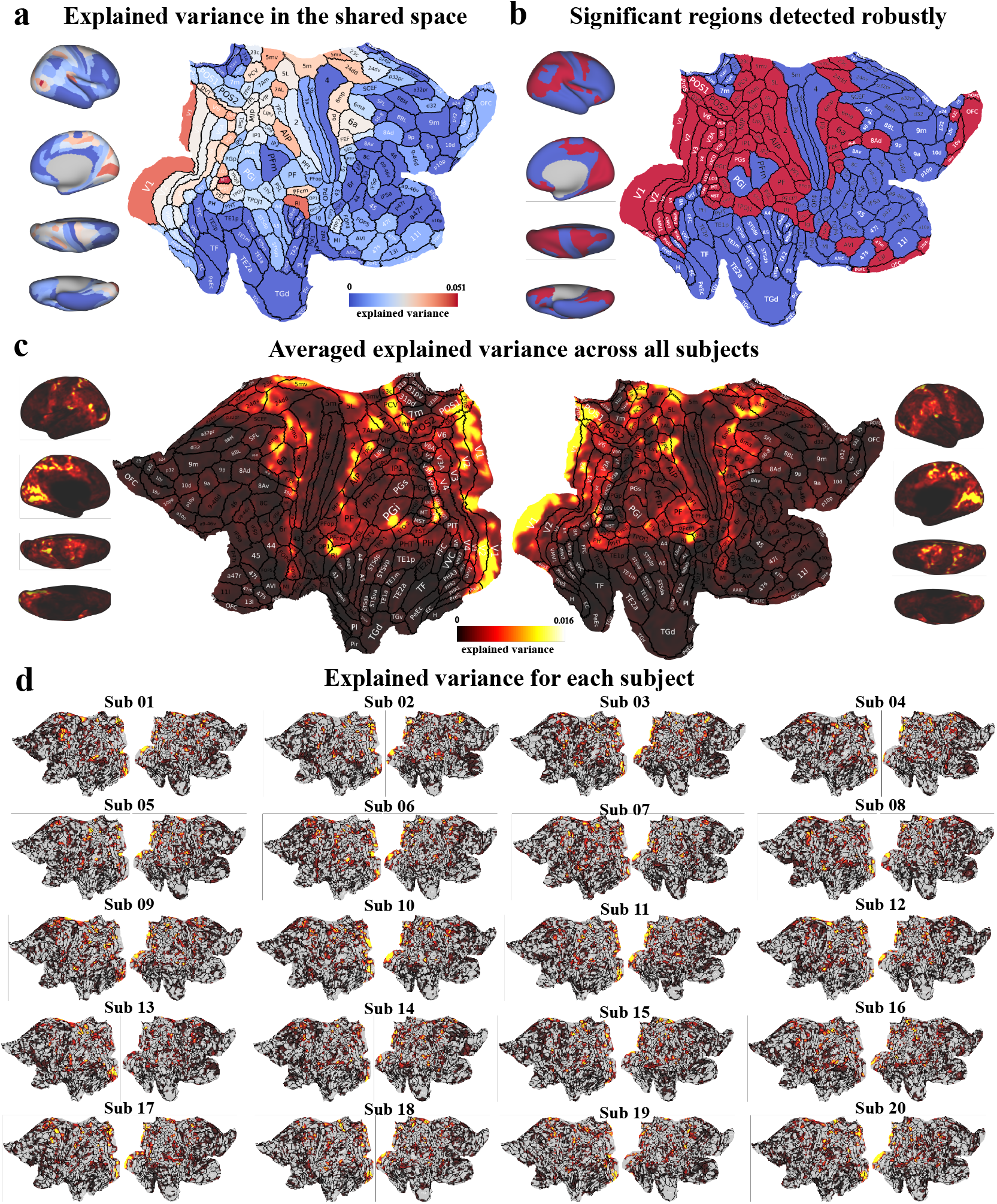
3D geometry representations across cortical regions. (**a**) **Explained variance for each region in the shared latent space of neural signals**. Colors reflect prediction performance by leave-one-run-out cross-validation. Red indicates more variance is explained. The sum of dimensions of shared temporal responses across the whole brain is 1409. (**b**) **Significant regions detected robustly**. To overcome the influence of the selection of SRM dimension on the result, we ranked the number of regions determined as significantly explained by geometry representation (p=0.05, permutation test) over a wide range of total SRM dimensions across the whole brain. The map displays the regions that are consistently determined as significant among the top 20% ranked results. The significant regions can be grouped as early-to-middle-level visual areas: V1, V2, V3, V4, V6, V7, V8; dorsal pathway: intraparietal cortex (AIP, VIP, LIPv, LIPd, MIP, IPS1, IP0, IP1, IP2), Brodmann area 7 (7PC, 7AL, 7Am, 7Pm, 7m, 7PL); medial pathway: MT+ complex regions (LO1, LO2, LO3, V4t, MT, MST, FST), temporo-parieto-occipital junction (TPOJ1, TPOJ2, TPOJ3), STV, PSL, parietal area F (PF, PFcm, PFt), RI; ventral pathway: ventromedial visual area (VMV1, VMV2, VMV3), parahippocampal area (PHA1, PHA2, PHA3), VVC, ProS, PreS, hippocampus; and regions serving other functions such as area 2, area 5 (5L, 5mv), and area 6 (6mp, 6ma, 6d, 6a) for somatosensory processing and movement, 23c and 24dd for behavior regulation, and OFC and pOFC for state representations. (**c**) **Averaged voxel-wise explained variance across all participants**. The shared response in testing data and prediction by encoding model were mapped back to the voxel space to calculate explained variance. (**d**) **Explained variance for each participant in voxel space**.

**Figure 4.**
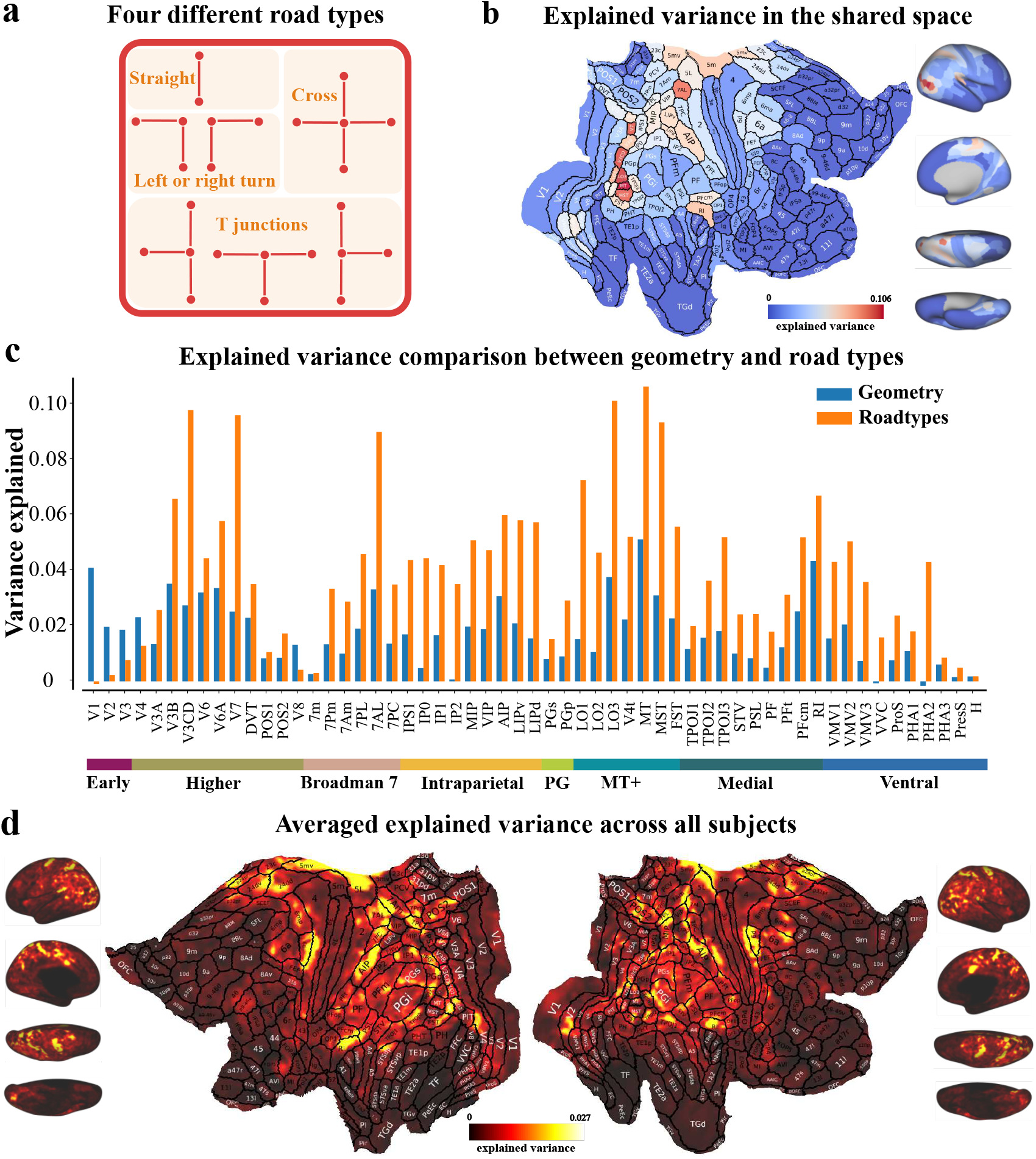
Abstract representations of road types across cortical regions. (**a**) **Four different road types**. The navigational routes were divided into segments of four types: unbranched straight road, left or right turns, T junction, and crossroads. (**b**) **Explained variance in the shared space**. Colors reflect prediction performance by leave-one-run-out cross-validation. (**c**) **Comparison of explained variance between compact representation of 3D geometry and road types across regions**. The explained variance of the early visual regions V1, V2 and V3 is much weaker for road types than for 3D geometry representation. (**d**) **Averaged voxel-wise explained variance across all participants**. It also shows that V1, V2, V3 are lower than other regions on the three pathways.

**Figure 5.**
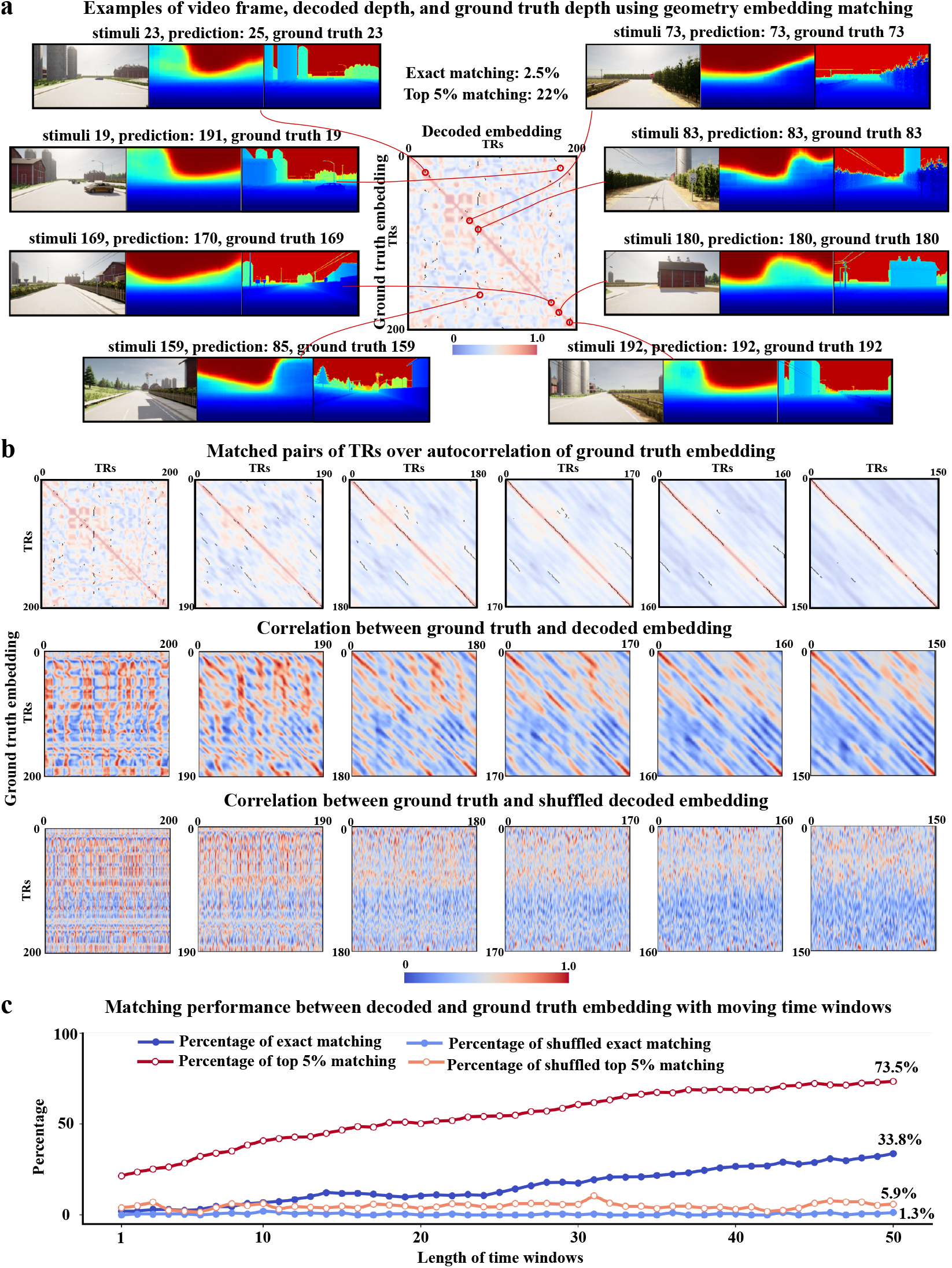
Reconstructing geometric structures. (**a**) **Examples of frames of stimuli, decoded depth, and ground truth depth images**. In the middle graph, the warmth of the background color indicates the correlation among the ground truth HRF-convolved latent vectors (embedding) of the deep generative model of depth across time. The black dots on the foreground show the TRs of the ground truth embedding (horizontal axis) best matching each decoded embedding (vertical axis). Left: examples of mismatch in geometry decoding. The decoded embeddings for TRs of 23 and 169 were mismatched to ground-truth embeddings of nearby TRs due to temporal autocorrelation. Those of TRs 19 and 159 were mismatched to embeddings far apart in time but with similar scene geometry structures. Right: examples of exact matching in geometry decoding. (**b**) **The effect of time window on correlational structures of embeddings**. As embeddings in longer time windows are included in calculating correlation, both the autocorrelation matrix of the ground-truth embedding (top) and the correlation matrix between ground-truth and decoded embeddings (middle) become more diagonal, while the the ground-truth and shuffled decoded embedding (bottom) keep uncorrelated. As a result, more black dots (best-matching TR-pairs) are located on the diagonal line with longer time window (top). Columns correspond to time windows of 1, 10, 20, 30, 40 and 50 TRs. (**c**) **The effect of increasing the time window during embedding matching**. As the time windows of decoded embedding to be matched with the ground-truth embedding increases to 50, the exact matching accuracy and the top 5% matching accuracy reach 33.8% and 73.5%, respectively, while the results using shuffled decoded embedding remain at chance level.

**Figure 6.**
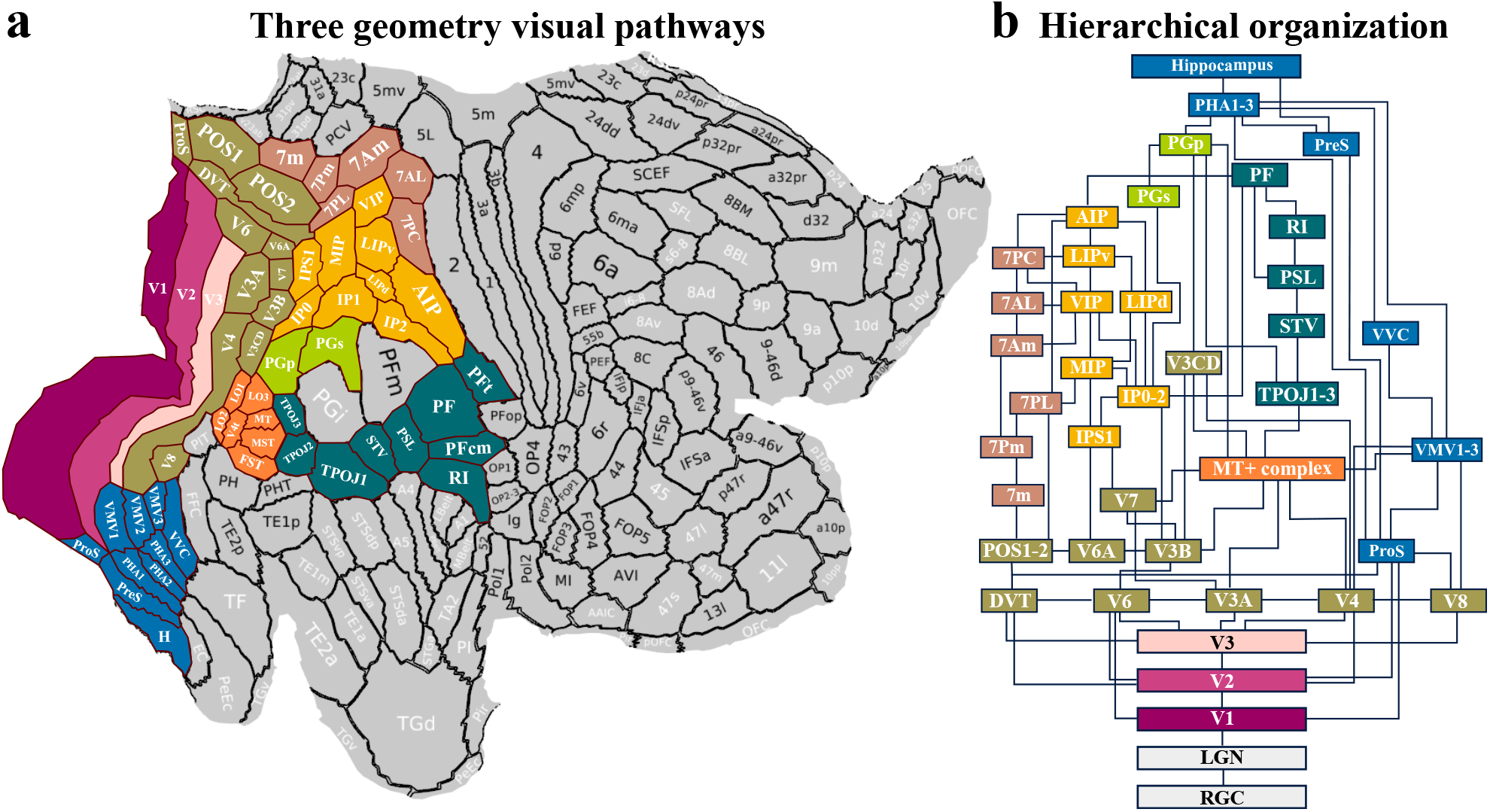
Illustration of three geometry visual pathways. (**a**) **Three geometry visual pathways on flattened brain surface**. The identified geometry-processing streams originate from the early visual cortex (V1, V2, V3), then reach the middle-level visual cortex (V4, V6, V7, V8), and branch into three different pathways: (1) the dorsal pathway reaches intraparietal areas (IP0, IP1, IP2, IPS1, MIP, LIP, VIP, API) and area 7 (7m, 7Am, 7Pm, 7AL, 7PL, 7PC); (2) the medial pathway first reaches the MT+ complex regions (FST, LO1, LO2, LO3, MST, MT, and V4t), then IPL areas (PFt, PF, PFcm) via TPOJ, STV, and PSL; (3) the ventral pathway reaches the parahippocampal gyrus PHA1-3 regions via VMV and VVC, then ProS and PreS, and finally arrives at the hippocampus. The dorsal and medial pathways merge together by the connections between IPL areas and IP area, which are separated by PFm and PGi. (**b**) **Hierarchical organization of the geometry visual system**. The diagram shows the connectivity according to the adjacency of regions on the brain surface. Colors code for different stages and pathways. The connectivity from PGp to PHA regions likely leads the dorsal and medial pathways finally to the hippocampus.

### The fMRI experiment of geometry representations

To identify brain regions carrying geometry representations, twenty participants watched videos of 3D scenes in eight different scenarios while undergoing fMRI scan (Figure 1). First, to eliminate the influence by low-level visual features, all participants watched videos with the same geometric structures, but with random weather and lighting conditions (Figure 1a). Second, to elicit proper brain activity of geometry representations in conditions close to real life, we used continuous, naturalistic movies taken from the driver’s point of view recorded while a car navigates in eight virtual towns with diverse geometric features as stimuli (generated by Carla [28], an open-source simulator for autonomous driving research, Figure 1b), instead of static scenes displayed one by one as in previous studies [18]. After watching each video for five minutes, participants responded to two memory tests of scenes in the video, to ensure their concentration on the video (Figure 1c).

### Modeling framework

We combined region-based shared response modeling (SRM) [26], a Bayesian algorithm of hyperalignment [27] implemented in BrainIAK [29, 30], and a deep generative model (a variational autoencoder) tasked to infer scene depth structure, to identify the brain regions that carry neural signals of geometry representations (Figure 2). To evaluate the contribution of every brain region to the representation of scene geometry, we separately applied SRM to each of 180 pairs of bilateral corresponding brain regions after parcellating the brains into 360 regions using HCP-MMP atlas [31]. We first estimated the optimal numbers of principal components [32] to explain the major signals of each region in each participant, and then used their average across participants as the maximum dimension needed to capture the synchronized signals for that region (Figure 2a). In parallel, we trained a deep generative model, which learns a few interpretable latent factors that capture the depth structures across a wider set of scenes from the virtual environments used in our study (Figure 2—figure supplement 1) and a mapping from the RGB images across weather and lighting conditions to these factors. We then built an fMRI encoding model to examine the amount of variance of the shared neural responses in each brain region explained by the geometric information captured by the latent factors of the deep generative model. We also trained a neural decoder to reconstruct depth structures by identifying the latent embeddings of the deep generative model for all scenes in a video that best match the latent embeddings linearly decoded from the shared neural responses (Figure 2b).

### Geometry representations over cortical regions

We examined the shared neural responses evoked by movies to identify the brain regions carrying the global geometric information captured by the deep generative model. Figure 3a shows the ratio of variance in each region explained by the geometric factors of scenes based on the trained encoding model using leave-one-run-out cross-validation. Regions in which the explained variance is statistically significant based on the permutation test are shown in Figure 3b (details in Figure 3—figure supplement 3). As the dimensions of the subspace of SRM influence the results, only the regions that are consistently detected among the dimensionalities of subspace that ranked in top 20% in terms of the total number of significant regions detected are considered as robustly detected regions (Figure 3—figure supplement 2). We further mapped each participant’s neural signals in the SRM latent space back to the whole-brain voxel space, and calculated the ratio of variance in voxel space that is explained by the encoding model. The averaged map of explained variance across participants (Figure 3c) shows a similar spatial pattern as those of both the region-based explained variance (Figure 3a) and the significant regions detected robustly (Figure 3b). Individual maps of explained variance are displayed in Figure 3d.

All different approaches of visualization show a consistent spatial pattern of three contiguous streams carrying geometry-related signals that start with the majority of the visual cortex (V1, V2, V3, V4, V6, V7, V8) located in the occipital lobe. Beyond these, the dorsal stream mainly includes the intraparietal cortex (AIP, VIP, LIPv, LIPd, MIP, IPS1, IP0, IP1, IP2), Brodmann area 7 (7PC, 7AL, 7Am, 7Pm, 7m, 7PL), and PG (PGs, PGp). The medial stream includes regions of the MT+ complex (LO1, LO2, LO3, V4t, MT, MST, FST), temporo-parieto-occipital junction (TPOJ1, TPOJ2, TPOJ3), STV, PSL, parietal area F (PF, PFcm, PFt) and RI. The ventral pathway includes the ventromedial visual area (VMV1, VMV2, VMV3), parahippocampal area (PHA1, PHA2, PHA3), VVC, ProS, PreS, and hippocampus. In addition to these geometry-encoding areas, a few regions traditionally indicated for other functions, such as area 2, area 5 (5L, 5mv), and area 6 (6mp, 6ma, 6d, 6a) for somatosensory processing and movement [33], 23c and 24dd for emotional and behavioral regulation [34], and OFC for representing reward value in decision-making or abstract task states [35, 36, 37] (abbreviations of regions are based on the Glasser atlas [31]).

### Abstract geometry representations in higher-level visual processing regions

We further investigated whether a more abstract representation of geometry beyond the depth structure, namely that of road structures, are carried in the same set of brain regions, using the same approach based on SRM. First, we annotated the type of road structure viewed at each TR in the movies as straight road, left or right turn, T junction, or crossroads (Figure 4a). Then, we fitted an encoding model for each region to predict the shared temporal responses based on the binary-coded time courses of road type convolved with a canonical hemodynamic response. Figure 4b shows the ratio of variance in each region explained by road types based on the trained encoding model using leave-one-out cross-validation.

Overall, the representation for the road types exhibits a spatial pattern of explained variance on the cortex similar to that for the pixel-wise depth. The early visual areas, V1, V2, and V3, are strongly involved in the geometry representations of pixel-wise depth structures, but less in in the abstract geometry representations of road types, which is demonstrated by the higher average voxelwise explained variance for depth-based geometric structure in these regions in Figure 4c. However, beyond these regions, the explained variance becomes larger for the abstract geometry representations of road types than for the depth-based geometric structure.

### Reconstructing geometric structures

We further demonstrate the possibility of reconstructing geometric structures by training a linear decoder that maps from the shared neural response signals to the time courses of the latent embeddings in the deep generative model. We averaged the shared neural responses in a left-out virtual city (Town07) over 7 repetitions experienced by each participant in different weather and lighting conditions as test data and then applied the decoder to them to reconstruct geometric structures. The latent embeddings of the scenes in the virtual town that best match the decoded embeddings are used to generate pixel-wise depth images with the deep network (Figure 2—figure supplement 1b). Examples of the matching result are shown in Figure 5a. Figure 5—video supplement 1 and 2 illustrate the RGB stimuli, decoded depth, and ground-truth depth images for time windows of lengths 1 and 50, respectively. As there is a strong temporal autocorrelation in the scene structure and similarity of depth structure between some scenes (shown by the background color of the middle subfigure of Figure 5a), scenes that are close in time or with similar geometry structure to the correct scenes are often mismatched (see the black dots off-diagonal in the middle subfigure of Figure 5a, and examples of mismatches on the left). Nonetheless, the correct scenes fall within the top 5% best-matching ones for 22% of the time. Further, identifying scenes by matching decoded embeddings in a longer time window improves the scene identification accuracy. As the time window increases to 50 TRs, the top 5% matching accuracy reaches 73.5% and the exact matching accuracy reaches 33.8%. As a control, the matching accuracy against shuffled true embeddings stays at chance level (Figure 5c).

## Discussion

It has long been proposed [38] that the visual system is composed of two separate streams of visual information processing, the dorsal (“where”) stream for spatial perception or action planning and the ventral (“what”) stream for object recognition and memory. The processing of scene geometric structure is key for spatial perception. However, a comprehensive characterization of the brain regions carrying global scene geometry information is lacking. By designing 3D outdoor scene stimuli that disentangle geometry structures from the variation in low-level visual features induced by weather and lighting conditions, this study identifies a fine-grained map of the cortical regions that participate in processing geometry information as shown on the flattened brain(Figure 3b, Figure 4b).

### Widespread distribution of representations for 3D scene geometry and abstract road types

Firstly, we identified a series of regions extending from V1 in the occipital lobe to intraparietal areas and Brodmann area 7 in the parietal lobe, consistent with the classical hypothesis of the dorsal “where” pathway specialized for representing spatial information [39]. However, we found that the geometry information of scenes is also represented in two additional medial and ventral pathways of contiguous regions. The medial pathway starts from the MT+ complex regions, and then reaches IPL areas. The ventral pathway reaches the parahippocampal gyrus PHA1-3 regions via VMV, VVC, PreS, and finally arrives at the hippocampus. Further, the representations of the 3D geometry of pixelwise depth structure and the abstract road structures are carried by overlapping regions along the three pathways, but the information of abstract road structures is absent in the early visual areas. We describe the three pathways jointly as “geometry visual pathways”. They appear wider than the regions identified to represent 3D surface in a previous study also using videos of diverse scene structures [17]. One possible cause is the difference in the representational model investigated. The previous study modeled 3D scene structures by the histogram of pixel-wise depth and surface norm, which discards the 2D-location information of each pixel and might map different geometry structures to the same histogram, while our deep network summarize major dimensions of variation of scene geometry structures.

The deep generative model allows the geometric structures of scenes to be reconstructed from the identified brain regions. It is worth noting that the synchronized neural signals evoked by the stimuli are of higher dimension than the compact representation of our deep generative model. This is because the deep generative model only captures the rough landscapes of the geometry using eight orthogonal units reflecting, e.g., e.g., blockage in the center, open field on the left or right, etc. When we perform reconstruction by matching predicted latent embeddings with ground truth latent embeddings in the network, mismatches occur at nearby time points or other scenes with similar global structure because the fine-grained geometry information is ignored by the networks Figure 5a. This can be overcome by lengthening the time window of embeddings to match.

### Illustration of geometry visual pathways

One can visually recognize from Figures 3a and 4b that the regions carrying geometry representations of the pixel-wise depth structure and abstract road type appear as three pathways starting from early visual cortex. We provide a diagram to illustrate the composition of the three-stream geometry pathways with connections based on spatial adjacency, and visualize them as a continuous map on the flattened brain in Figure 6a. The pathways start from early visual areas of V1, V2, and V3. A stripe of mid-level processing areas connects these early visual areas with the separate dorsal, medial, and ventral pathways. The mid-level areas include V4, V3A, V3B, V3CD, V6, V6A, V7, DVT, POS1, POS2, and V8. Departing from the mid-level processing areas, the dorsal pathway reaches intraparietal areas (IP0, IP1, IP2, IPS1, MIP, LIP, VIP, AIP) and area 7 (7m, 7Am, 7Pm, 7AL, 7PL, 7PC). The medial pathway starts from the MT+ complex regions (FST, LO1, LO2, LO3, MST, MT, and V4t), and reaches IPL areas (PFt, PF, PFcm) via TPOJ, STV, and PSL. The ventral pathway first reaches VMV and VVC, then ProS and PreS, and through parahippocampal gyrus PHA1-3 regions finally arrives at the hippocampus.

The dorsal and medial pathways are separated by PFm and PGi, and merge together by the connections between IPL areas (PF, PFt) and IP areas (AIP, IP2) (Figure 6a). As geometry information should ultimately be utilized by the hippocampus, we speculate that the information from both dorsal and ventral pathways are likely fused and sent into the hippocampus via PHA regions by the PGp area due to its strong connectivity to PHA [40].

### Hypothetical roles of the three geometry visual pathways

We postulate different functions of geometry representations may arise through the three pathways and their interactions. The dorsal pathway forms a representation for global 3D surface [22, 41]. The medial pathway integrates multisensory information and depth cues including motion signals [42, 43], to distinguish objects and infer their shapes. The ventral pathway fuses the information from dorsal and medial pathways to represent objects, places, spatial layouts, and landscapes at the scene level for constructing cognitive maps [44]. These roles remain tentative and call for causal and active-navigation studies.

To elaborate further, the early visual areas (V1, V2, V3) extract low-level features such as edge, local motion, object contour and texture that support a wide range of visual inference [45]. The mid-level visual areas (V4, V3A, V3B, V3CD, V6, V6A, V7, DVT, POS1, POS2, and V8) might extract various cues of depth variation using the low-level features provided by the previous stage.

The dorsal pathway is considered here as “the geometry formation pathway”, mainly including the intraparietal areas (IP) and area 7, which estimate environmental geometry (3D surface structure) from cues provided by earlier stages [46]. This geometry information is further used for the guidance of movements of reaching, grasping, and manipulating objects, spatial attention, and navigation carried out in IP1, MIP, LIP, VIP, AIP, 7Am, and 7Pm [47, 48]. Some areas, such as IP0-2 and 7m, are responsible for representing object location and relationships between objects [49]. Regions like LIP and 7AL direct spatial attention to focus on important information [50], while spatial working memory (carried by IPS1) further helps to merge the 3D geometry structure inferred from different viewpoints to form a local geometric map [51].

We consider the medial pathway as “the shape inference pathway”, mainly including the MT+ complex regions (FST, LO1, LO2, LO3, MST, MT, V4t), TPOJ, STV, PSL, and IPL areas (PFt, PF, PFcm). The MT+ complex regions estimates the shape and structure of objects from visual motion, such as optical flow [52]. The integration of multisensory information, including visual, auditory, and tactile inputs in regions such as TPOJ, STV, PSL, and IPL areas (PFt, PF, PFcm) [53] could further facilitate shape inference. For example, tactile information from the invisible back side of an object can augment the surface geometry provided by the dorsal pathway to help complete the estimation of its shape. The medial pathway may also integrate optical flow information provided by MT and MST [54, 55] for inferring distances of objects.

The ventral pathway is considered as “the scene representation pathway” [40]. ProS, VMV, VVC, and PreS are mainly involved in object recognition and motion perception [56]. PHA1-3 are associated with representation of scenes, such as landscapes, indoor environments, and spatial layouts, which help individuals to understand their surroundings for spatial navigation [19]. The holistic geometric shape and structure representations formed by the medial and dorsal pathways are likely integrated with the local texture information extracted in the “what” visual pathway to support semantic inference [57]. This allows the brain to distinguish similar textures even in chameleon camouflage, dynamic traffic, and pedestrian flow in different weather and lighting conditions.

Under this hypothesis, the three pathways are not independent. The representation of the 3D geometry of visible surfaces might be first generated by the dorsal geometry pathway and then broadcast to both the medial and ventral pathways to support inferring holistic shapes and scene layouts, which explains the wide distribution of 3D geometry representation. Why then is the more abstract road type information also carried by all three pathways? This is compatible with the view that different parts of the visual system jointly perform probabilistic inference over a hierarchical generative model of visual scenes, composed of nodes of latent variables and their causal relations [24, 25]. In this view, a latent variable represented in one region constrains both its child nodes as a prior and parent nodes through a learned likelihood function, which requires reciprocally sending signals between regions [58]. Road type information may be formed in the ventral pathway but fed to the other two pathways to provide such constraints.

### Methodological considerations

Participants were not required to fixate in our study to make the experience resemble natural real-world navigation. Hyperalignment has been demonstrated to generalize the function-to-brain mapping across participants precisely without requiring fixation [59].

One unanswered question is the role of optic flow in geometry representation. Because optic flow is intrinsic to naturalistic navigation videos, activity in MT/MST likely reflects a combination of motion sensitivity and geometry perception. Importantly, optic flow itself is a fundamental depth cue: the global pattern of motion across the retina directly depends on the 3D structure of the scene [60, 61]. Thus, our results should be interpreted as reflecting a combination of geometry- and motion-related signals. However, neural responses to optical flow alone are not sufficient to explain our findings, as the results demonstrate geometry and affordance representations extend well beyond motion-selective cortex into distributed dorsal, medial, and ventral pathways.

### Challenges for understanding geometry representations in the brain

Very few studies have attempted to bridge the research on visual processing in the cortex and that on hippocampal cognitive maps for spatial navigation in the limbic system [62]. Application of deep networks to model brain activity has primarily focused on the “what” aspect of vision [63, 64], while the representation of geometry is perhaps more directly linked to cognitive maps. The representations of 3D scene geometry and road types arising in the visual pathways support higher-level representation of the spatial layouts of one’s surroundings [16, 15] through boundary representations [11]. The layout representations join semantic representations along the “what” visual pathway together to support neural representations of animals’ positions and headings by grid cells, place cells and head direction cells [65]. By focusing on the geometry representations critical for spatial layouts, our research takes one step towards bridging the fields of vision and cognitive map. Yet, it is still challenging to understand the computational mechanisms by which the geometry representations transform into those of the hippocampus step by step for complex navigation. Future works will benefit from exploring both explainable models [17] and deep networks [66] in search for the representations used by the brain throughout the transformation, and involving active navigation of participants in virtual environments with natural scenes [67].

While our grouping of the regions into three pathways is based on their spatial adjacency on the HCP-MMP atlas [31], many regions are connected by long-range axonal projections [68]. A major challenge is to further understand the interactions within and among the dorsal, medial, and ventral pathways for geometry representations. Future causal studies that intervene (e.g., by TMS) regions in the three pathways and examine the impact on geometry representations might prove fruitful.

## Methods

### Experimental paradigm

The study was approved by the Office for Life Science Research and Ethics, The University of Tokyo. After giving informed consent and screening for MRI safety, each participant passively watched 14 videos recorded from the first-person driver’s view of a vehicle moving in 8 different virtual towns with pre-planned routes while their brain activities were scanned by fMRI. One of the towns was experienced for 7 times, each with different weather and lighting condition. Participants were instructed to focus and enjoy watching the movie. To ensure attention, after each video, they were presented with two images, one from the video they had just watched, the other taken from another town under the same weather condition, and were asked to judge which one they had just seen in the movie. A full-brain structural MRI image was acquired before the video-watching task, and a pair of spin-echo images were acquired after half of the fMRI session. After exiting the scanner, participants rated their sleepiness, tiredness, the degree of feeling immersed in the virtual reality environment, and what came to their mind during the experiment (not analyzed here).

### Participants

20 participants took part in the study, 15 males and 5 females. Participants’ ages range from 19 to 37 (M:27.3; SD:6.0). 19 were Asian and 1 was Caucasian. Except for 2 participants who were authors of the paper, all other participants were naive about the purpose of the study.

### MRI data collection

All MRI data were collected with a Siemens 3T Prisma MRI scanner at International Research Center for Neurointelligence, The University of Tokyo, with a 32-channel head coil. Each participant completed all the scans in one session, including a T1-weighted structural scan at 0.8mm resolution and 14 gradient-echo multi-echo echo-planar imaging (EPI) scans for acquiring fMRI data of the main task. The EPI data were acquired with multi-band sequence produced by Center for Magnetic Resonance Research (CMRR) of University of Minnesota, with a homogeneous voxel size of 2.8 × 2.8 × 2.8 *mm*^3^, whole-brain coverage by 45 axial slices each with 70 × 70 voxels. Both phase and amplitude images were recorded at five echo times (TE): 14.0, 41.51, 69.02, 96.53 and 124.04 ms (only amplitude images were analyzed) and at a repetition time (TR) of 1.5 s with a flip angle of 66°and echo spacing of 0.496 ms. A multi-band acceleration factor of 5 was used, together with a 75% partial Fourier in-plane acceleration. Spin-echo images were recorded with the same and opposite phase encoding directions as the EPI data, both with identical resolution, brain coverage, bandwidth, echo spacing, and slice placement as the EPI data for the purpose of correcting in-plane distortion due to the inhomogeneity of magnetic field. The TR was 8000 ms and TE was 66 ms for the spin-echo data.

### Stimuli

In order to elicit representation for 3D structures of commonly seen visual scenes, we generated video recordings of the scenes observed from a front-facing camera on a virtual vehicle moving in 8 different towns with diverse types of landscapes (e.g., rural, urban, highway, etc.), rendered with an open-source simulator for autonomous driving research, Carla. The vehicle moved at a speed of approximately **30** miles/hour following a predefined route. Example views in the towns and bird-eye views of the vehicle’s routes are displayed in Figure 1—figure supplement 4. In order to disentangle the representation of geometry from that of low-level visual features such as brightness, edges of shades, and reflection, we generated multiple videos recorded under different weather conditions or time in a day for each town, following the same navigation path. The weather and time of recording for each town’s video being watched were randomly assigned to different participants, except for one town from which participants all watched 7 videos with different weather and time of day following the same path.

### Learning compact latent representations of 3D scene geometry

To search for a compact, low-dimensional representation of 3D geometry commonly observed in outdoor environments, we trained a deep neural network that processes 2D images and represents their 3D depth structures with a low-dimensional vector. We adapted the beta-variational auto-encoder [69, 70] that has the strength of learning a disentangled representational space such that traversing in it allows smooth interpolation among data samples and that each dimension captures meaningful and distinct variation in data. The neural network is composed of an encoder that infers the mean and standard deviations of a diagonal covariance matrix of the posterior distribution of the latent vector for each input 2D image. Samples drawn from the posterior distribution are then passed through a decoder to predict the logarithm of depth in each pixel. Both the prediction error for the logarithm of depth and the Kullback-Leibler divergence between the posterior distribution and a standard multi-variate normal distribution are optimized to train the networks. We used Vision Transformer as the architecture for both the encoder and decoder. To train this model, we used much more diverse visual inputs taken from many more routes in the towns used for the study, which includes scenes not seen by participants during the experiment.

### fMRI data preprocessing

fMRI data were preprocessed primarily using `afni proc.py` command in AFNI [71] (version 22.1.06) on an Ubuntu 20.04 operating system. After slice-timing correction of the EPI data, spatial distortion along the phase encoding direction was estimated using the spin-echo images with opposite phase encoding directions, and the correction for this distortion was applied to the EPI data. The resulting EPI data were then corrected for head motion, and the anatomical image of the brain (after stripping the skull) was aligned to the EPI data by rigid-body transformation only. The EPI data acquired at 5 different echo times were then weighted voxel-wise with an optimal weighting procedure [72, 73]. The intersection of the whole-brain mask derived from the aligned anatomical image and combined EPI images was used to select voxels within the brain.

In parallel, a nonlinear spatial warping was estimated to align each participant’s anatomical image with the MNI template (`MNI152_T1_2009c_in_AFNI`). To perform analysis within each region, the volumetric HCP-MMP (Glasser) atlas [31] in MNI space was inverse-warped to each participant’s anatomy using AFNI’s nonlinear transformation [71, 74].

### Shared response model

After temporal z-scoring of fMRI time courses within each run, the shared response model (SRM) was fitted to the time courses within voxels in each of the brain regions in the Glasser atlas. SRM takes fMRI time courses from a selected region of each participant who experience a synchronized task (in our case, watching videos recorded from the same driving route in the same town but with different weather/time of day across participants), and simultaneously estimates a set of time courses shared across all participants and a set of individualized orthonormal spatial weight matrices for each participant that project the time courses to the space spanned by all voxels as axes to maximally explain the variance of the fMRI data[26]. Because the variation of neural signals introduced by different lighting conditions is not synchronized across participants, this approach will maximize the chance of identifying neural signals corresponding to only the synchronized representations of the visual input, including the variation of geometric properties of scenes. The learned spatial weights can be used to project left-out data to the shared latent space in any cross-validation procedure to evaluate encoding or decoding models.

### Road type annotation

A research assistant annotated the type of road structure at the current location of the viewer in the video into four categories for the frame at each TR (which was used to calculate embedding for depth structure as well). The four categories are: continuous road without branching (this includes both straight roads and when a road winds), left or right turn, being at a T-junction, and being at a four-way cross.

### Encoding model and explained variance

For both the compact representation of scene geometry in our fitted deep network and the abstract road type representation, we used k-fold (*k* = 7) cross-validation approach to estimate the proportion of variance in the shared neural response across participants that can be accounted for by each representation separately. More specifically, in each fold, SRM was fitted to z-scored fMRI signals in each parcel (region) *m* of the Glasser atlas [31] of all participants in 6 of the 7 runs in which participants watched a video navigating in a town only once. In brief, SRM estimates orthonormal spatial weights 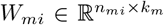 (*n*_*mi*_ is the number of voxels in region *m* of participant *i* and *k*_*m*_ is the dimensionality of the shared response model for that region) for each participant *i* such that the neural signals 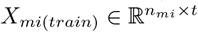 (*t* is the total time points) in the corresponding region of the training data are best reconstructed by multiplying the spatial weights with a common set of time courses 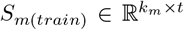 in a latent space shared across participants that are simultaneously estimated: *X*_*mi*(*train*)_ = *W*_*mi*_*S*_*m*(*train*)_ + *ϵ*_*mi*(*train*)_, where *ϵ*_*mi*(*train*)_ is residual to be minimized [26]. Spatial weights of all participants are all jointly estimated. We then used kernel ridge regression implemented in the Himalaya package [75] to fit linear encoding models *S*_*m*(*train*)_ = *P*_*m*_*Y*_*train*_ + *E*_*m*(*train*)_ that separately predict the shared neural signals based on the representation of 3D geometry or road types, using the re-gressors *Y*_*train*_ ℝ ^*h* × *t*^ (*h* is 9 for 8-dimensional time courses of the latent codes in the deep generative model or 5 for 4-dimensional binary codes of road types, both convolved with a canonical hemodynamic response function (HRF), padded with an additional offset component of the constant 1). A regularization term on the L2 norm of the regression coefficients *P*_*m*_ was included in the loss function in addition to the MSE loss of regression residuals *E*_*m*(*train*)_, and the weight of the regularization was determined by nested cross-validation. The estimated weights were then multiplied with the regressors for the testing run to predict the shared response in those runs. The explained variance was calcu-lated as 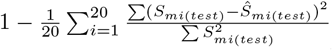, where 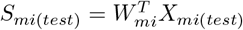 is the shared response time course of region *m* of participant *i* in the test data and *Ŝ* = *P*_*m*_*Y*_*test*_ is the predicted shared response based on the learned spatial weights and regressors. We rotated the choice of testing runs over the fMRI runs of watching videos from the 7 towns that were not repeated within participants, and calculated the average ratio of explained variance across all 7 folds.

To calculate voxel-wise explained variance, we first projected each participant’s shared response signal back to their brain using the spatial weight matrix *W*_*mi*_ learnt for each participant as 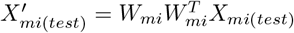, and then projected the shared response predicted by the encoding model also to the voxel space as 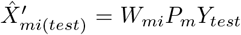. Then the ratio of explained variance is calculated per voxel between the signal 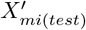 reconstructed by SRM and that predicted by the encoding model, 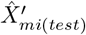.

To determine if the ratio of explained variance is statistically significant, we performed permutation test. In each permutation, we shuffled the time indices of the shared neural responses within each training and testing run, and then calculated the explained variance with the same procedure as for the original data. The p-value for each region is calculated by the frequency that ratio of explained variance in permuted data exceeds the ratio in the original data, out of 10,000 permutations. Then all p-values were corrected for multiple test using the Python module ‘statsmodels.stats.multitest.multipletests’. and regions with p-value smaller than 0.05 were considered significant.

We empirically found that the choice of the dimensions of the shared signals in SRM influences the variance of the shared response explained by the representation of geometry in our model, and consequently the number of significant regions. The largest number of regions is detected when in total 1409 dimensions are retained by SRM across regions (Figure 3—figure supplement 2a), based on which we display the explained variance across regions in Figure 3a. To reliably identify the fine-grained map of geometry representation, we evaluated 100 approximately equally spaced values of total dimensions of shared responses across all regions. To select the 100 dimensionalities, the sum of the optimal numbers of principal components (PC) [32], needed to capture major covariance structure in each region’s original signals in the voxel space averaged over participants, was chosen as the highest (5465). The lowest total dimension (225) was chosen by first scaling down the highest total dimension to the total number of ROIs (180) while keeping the ratios among SRM dimensions, and then rounding the resulting dimensions of each region to the nearest integers. As this resulted in zero dimensions for some regions, these zeros were replaced by one, which resulted in 225 total SRM dimensions across regions. All other evaluated total dimensions are approximately equally spaced between the highest and lowest total dimensions while keeping the ratios among the SRM dimensions across regions. To ensure the robustness of our finding, we first ranked the total SRM dimensions based on the number of significant regions detected, and then evaluated the regions that are consistently detected by the top-ranking total SRM dimensions. As shown in Figure 3—figure supplement 2c, as a wider range of top-ranking SRM dimensions are considered, the sets of significant regions are largely stable from considering the top-5 ranked dimensions to the top-60, except that TPOJ2 drops from the set of consistently detected regions after expanding to the top-30 ranked dimensions. Only OFC drops after including more top ranked dimensions from 70 until including all evaluated dimensions (100). We reported the regions consistently detected across all SRM dimensions that ranked in the top 20 in Figure 3b.

### Geometry decoding models

The decoding pipeline of the scene geometry structure shares some similarities as the encoding pipeline. We also used the Himalaya package to learn a linear regression model that uses the average shared responses to predict the time courses of the embedding of the depth generative network convolved with the canonical HRF. The shared responses were from all the regions detected as significantly explained by the deep network’s embeddings when the total number of SRM dimensionalities (1409) yielded the maximal number of statistically significant regions within the training data. After fitting the model to the fMRI data of the 7 towns without repeated viewing, we used the estimated weights to decode the time courses of the HRF-convolved embedding in the depth neural network for the 8^*th*^ town based on the average shared responses across all 7 repeated runs (under different weather and time of day) of all participants. Correlation between the decoded embedding and the embeddings of all TRs within the movie was calculated. The TR of which the ground-truth embedding had the highest correlation with the decoded embedding is treated as the decoded scene.

To overcome the false recognition due to temporal autocorrelation in the ground-truth HRF-convolved embedding, we also concatenated the decoded embedding within a time window centered at the time point of query and calculated their correlation with the concatenated ground-truth HRF-convolved embeddings within time windows of the same length, shifted across the entire movie, to decode the scene structure. This yields improved scene geometry reconstruction accuracy (Figure 5c).

## Data availability

The dataset is available on OpenNeuro in BIDS format at https://openneuro.org/datasets/ds005354. It includes both anatomical brain images and functional activity data.

## Code availability

The custom code used for conducting the experiments and analyzing the fMRI data can be accessed at https://github.com/geometry-in-human-navigation.

## Acknowledgement

The study was funded by WPI-IRCN startup budgets (M.B. Cai and T. Zeng) and JSPS KAKENHI Grant Number JP21K20679 (T. Zeng). The research does not reflect the funders’ opinions. We thank Aaron Tsubasa Nakamura for his tremendous effort in assisting the fMRI experiment and translating experimental materials. We appreciate the support of the WPI-IRCN Human MRI Core and Dr. Naohiro Okada, the University of Tokyo Institutes for Advanced Studies.

## Additional information

**Figure 5—video supplement 1** Illustration of the RGB stimuli, decoded depth, and ground-truth depth images for a time window of length 1.

**Figure 5—video supplement 2** Illustration of the RGB stimuli, decoded depth, and ground-truth depth images for a time window of length 50.

---

**figure supplement 1.**
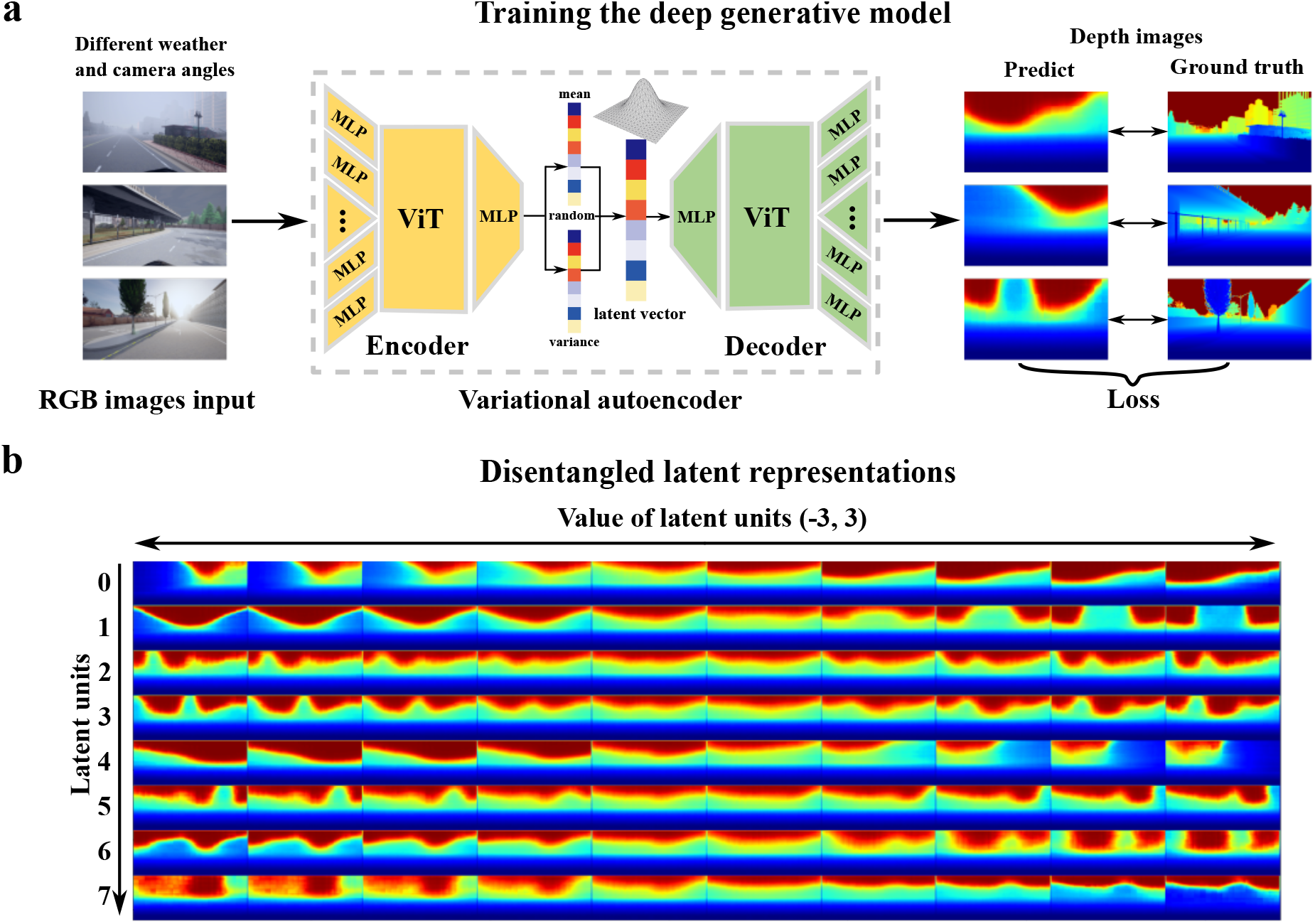
The deep generative model for learning latent representations of scene geometry. (**a**) **Model architecture inspired by beta-variational auto-encoder** Its encoder learned latent representations and their uncertainty (through standard deviations of a diagonal covariance matrix of the mean latent code) for RGB images. The images to train the network were taken by a camera with random yaw angles from −60^°^ to 60^°^ on a virtual vehicle navigating in the same set of virtual environments under different weather and lighting conditions, but through more diverse routes than those used to generate experimental stimuli. The depth images were generated by passing samples drawn from a multivariate normal distribution centered at the latent representations through the decoder. (**b**) **The disentangled representations**. Visualization of the variation in geometry structure encoded by each latent dimension of the deep generative model. In each row, the value of a single latent unit was set from −3 to 3 with equal spacing, while other units were fixed at 0. The results show that the units 0 and 4 both encode for views with close distance on one side of the scene, but in opposite ways. Unit 1 encodes for variation from roads with occluding structures on both sides to scenes in which the central view is blocked. Units 6 and 7 prefer lower buildings, while units 0 and 4 prefer higher buildings. Units 2, 3 and 5 prefer scenes with detailed landscape variation far away from the viewer.

**figure supplement 2.**
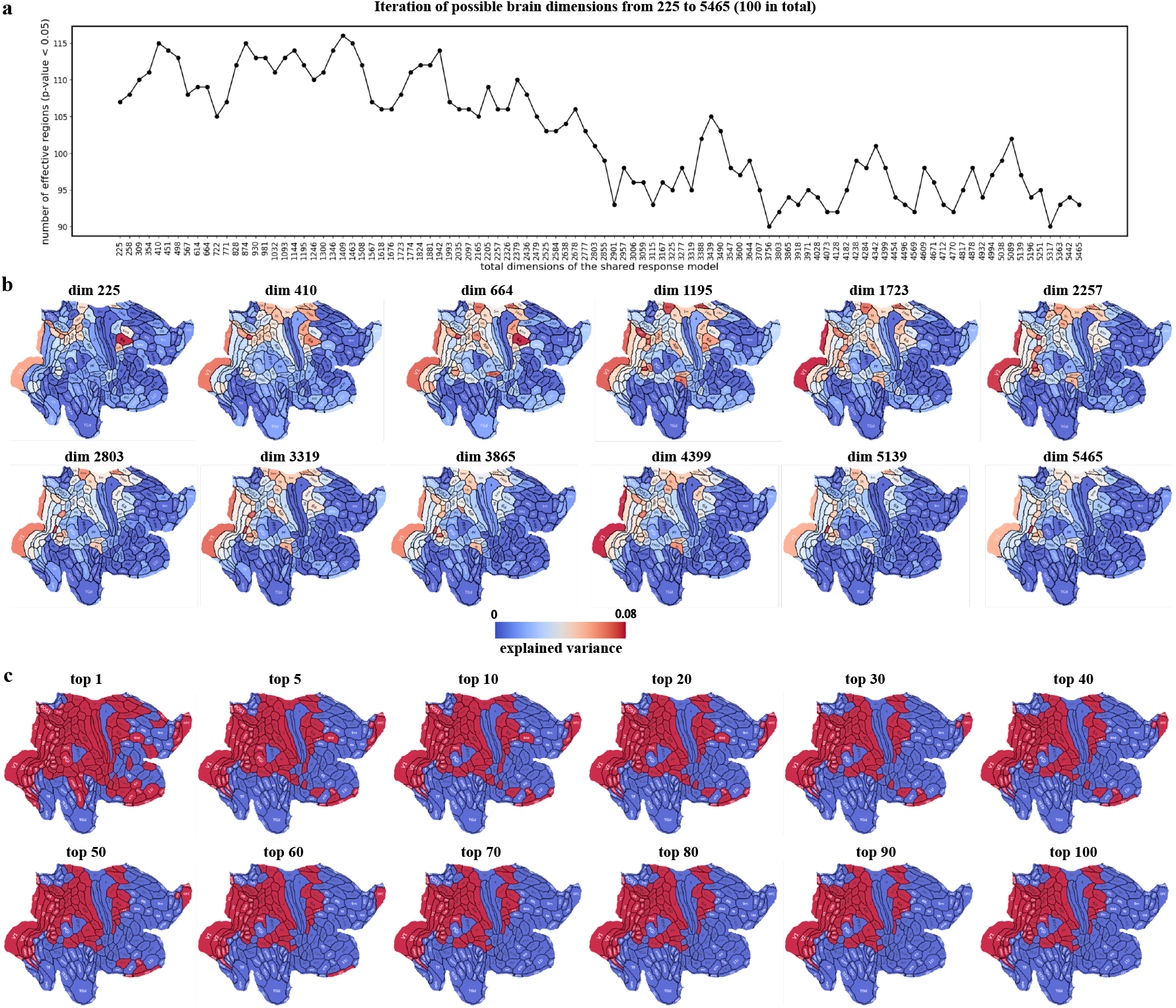
Procedure of identifying regions that can be robustly detected as encoding geometry information insensitive to the choice of SRM dimensions. (**a**) **Number of regions detected as significant when choosing different total dimensions of shared responses**. To determine the proper total dimension of shared responses across the whole brain, equidistant search was performed in 100 values from 225 to 5465 (refer to Method for the choice of the highest and lowest dimensions). The vertical axis indicates the number of significant brain areas detected. (**b**) **Examples of region-wise explained variance at different total dimensions**. Different colors indicate the levels of explained variance. (**c**) **Robustness in identification of significant regions**. We first ranked the total SRM dimensions based on the number of significant regions detected. The subfigure “top 1” shows the map of the largest number of regions detected as being significantly explained by geometry information across all choices of total dimension of shared response (it occurs at 1409 total dimensions). The subfigure “top 100” shows the regions that are always detected as significant across all 100 choices of total dimensions. Although from top 1 to top 100, the number of regions robustly detected as significant decreases, the sets of significant regions are largely stable from top 5 to top 60 ranked dimension. The same set of regions are detected by the top 70, 80, 90, and 100 ranked dimensions. 2

**figure supplement 3.**
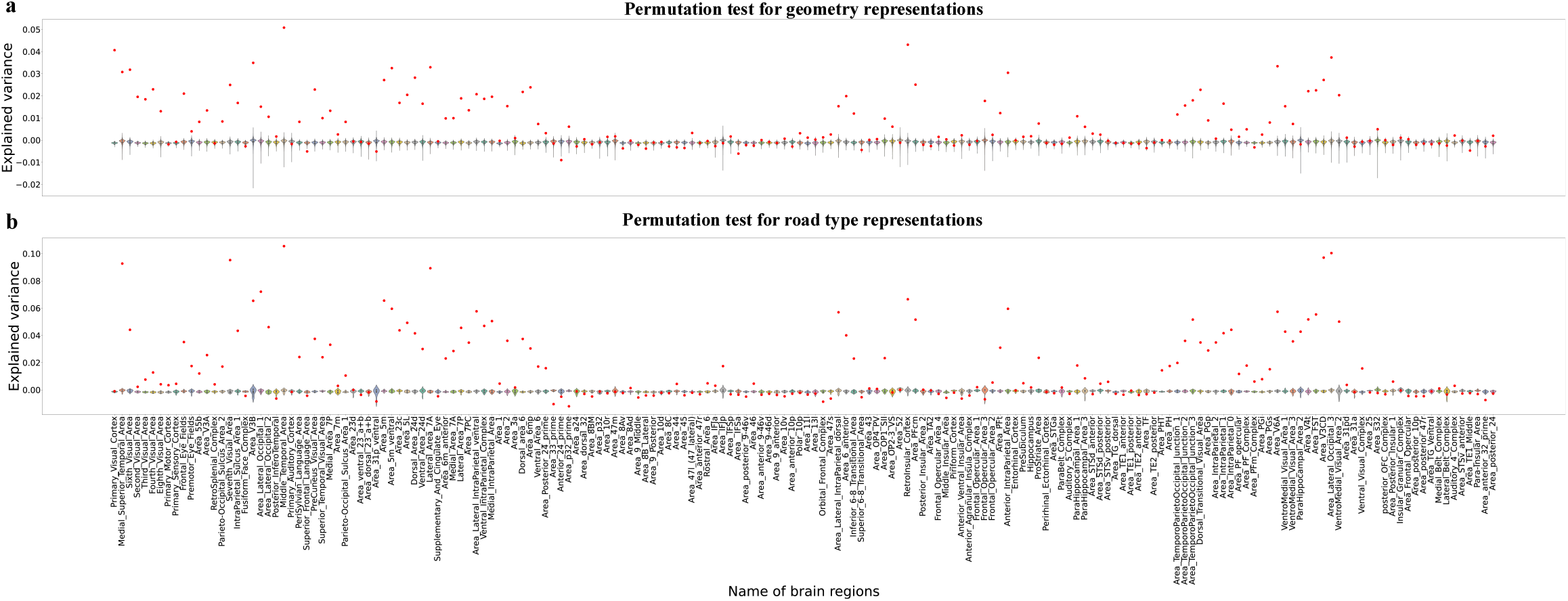
Permutation test results for **(a) geometry representations** and **(b) road type representations** across different brain regions. The red dots indicate the explained variance ratio obtained from the SRM data with 1409 total dimensions, while the violin plots represent the distribution of explained variance ratio obtained from 10,000 permutations that shuffle the TRs of the shared response signals in each run. Y-axis: ratio of variance in shared neural responses of each region explained by regressors of the corresponding representation. X-axis: brain regions.

**figure supplement 4.**
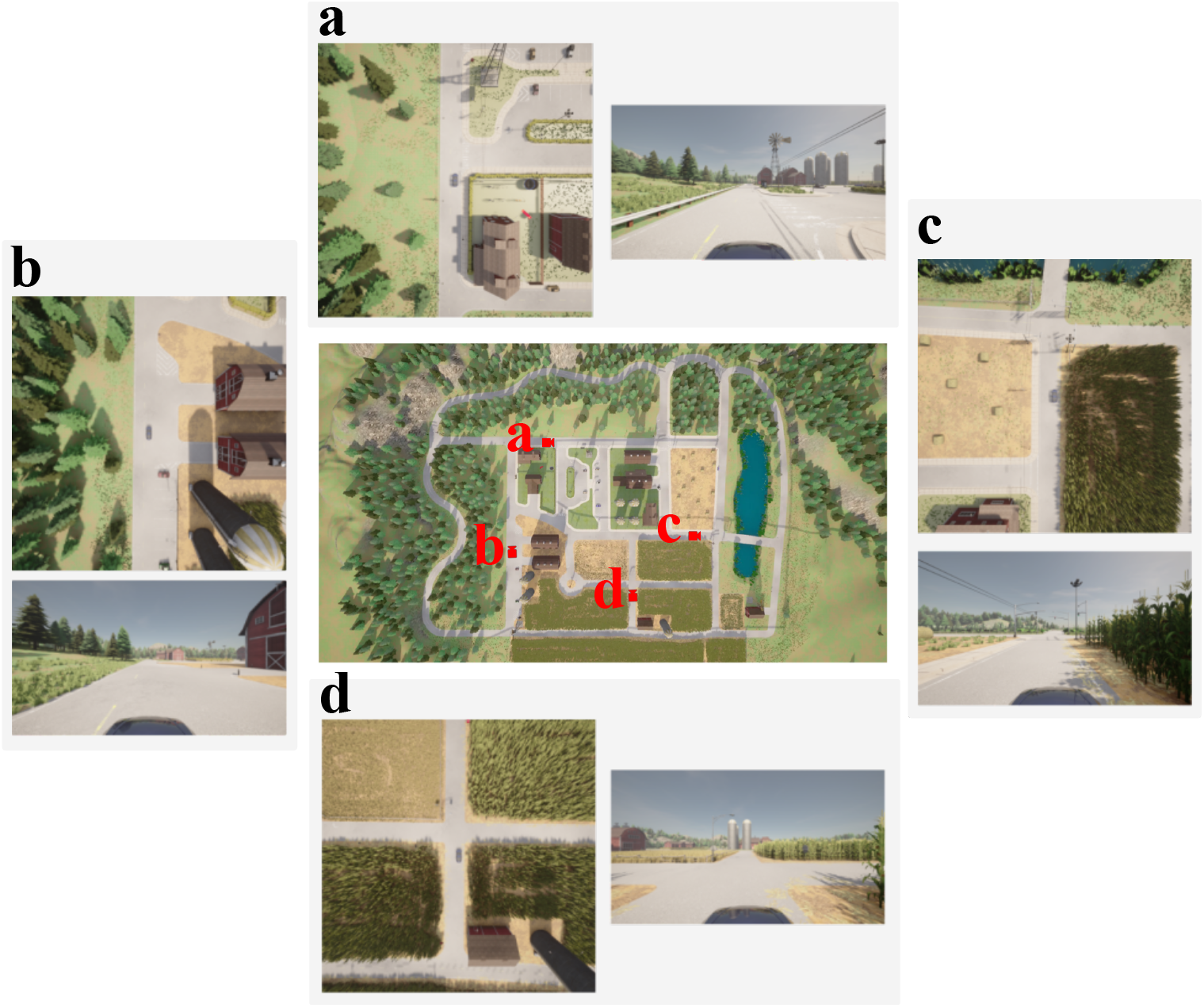
Examples of first-person views and bird-eye views at different locations along the vehicle’s routes in Town 07. The central image depicts the overall map of the environment with the locations marked by red dots **(a-d)**. Each marked location is accompanied by two images: a bird’s-eye view of local environment centered at the vehicle and a first-person perspective view from the vehicle.

